# Distinct Tumor-TAM Interactions in IDH-Stratified Glioma Microenvironments unveiled by Single-Cell and Spatial Transcriptomics

**DOI:** 10.1101/2024.05.23.595505

**Authors:** Meysam Motevasseli, Maryam Darvishi, Alireza Khoshnevisan, Mehdi Zeinalizadeh, Hiva Saffar, Shiva Bayat, Ali Najafi, Mohammad Javad Abbaspour, Ali Mamivand, Susan B. Olson, Mina Tabrizi

**Author notes:** Correspondence: Mina Tabrizi M.D., Ph.D. Meysam Motevasseli and Maryam Darvishi have equally contributed to this work.

## Abstract

Tumor-associated macrophages (TAMs) residing in the tumor microenvironment (TME) are characterized by their pivotal roles in tumor progression, antitumor immunity, and TME remodeling. However, a thorough comparative characterization of tumor-TAM crosstalk across IDH-defined categories of glioma remains elusive. We delineated the phenotypic heterogeneity of TAMs across IDH-stratified gliomas. Notably, two TAM subsets with a mesenchymal phenotype were enriched in IDH-WT glioblastoma (GBM) and correlated with poorer patient survival and reduced response to anti-PD1 immune checkpoint inhibitor (ICI). We proposed SLAMF9 receptor as a potential therapeutic target. Inference of gene regulatory networks identified PPARG, ELK1, and MXI1 as master transcription factors of mesenchymal BMD-TAMs. Analyses of reciprocal tumor-TAM interactions, revealed distinct crosstalk in IDH-WT tumors, including ANXA1-FPR1/3, FN1-ITGAVB1, VEGFA-NRP1, and TNFSF12-TNFRSF12A. Furthermore, we demonstrated significant upregulation of *ANXA1, FN1, NRP1*, and *TNFRSF12A* genes in IDH-WT tumors using bulk RNA-seq and RT-qPCR. Longitudinal expression analysis of candidate genes revealed no difference between primary and recurrent tumors. Collectively, our study offers insights into the unique cellular composition and communication of TAMs in glioma TME, revealing novel vulnerabilities for therapeutic interventions in IDH-WT GBM.

## INTRODUCTION

Glioma, the most aggressive and common primary brain tumor, constitutes approximately 80% of all brain malignancies, including glioblastoma (isocitrate dehydrogenase (IDH)-wild type (WT)) (GBM) and IDH-mutant (Mut) gliomas [1]. GBM is categorized as an incurable high-grade diffuse glioma, and the most significant challenge leading to therapeutic failure lies in its intra-tumoral heterogeneity [2, 3]. Among the four cellular states of GBM, with the exception of the mesenchymal-like (MES-like) state, the rest exhibit gene expression patterns that resemble those of normal neurodevelopmental cell types. Additionally, genetic drivers of GBM, such as alterations in *CDK4, PDGFRA*, and *EGFR* genes, play a crucial role in governing the emergence and prevalence of the neural progenitor-like (NPC-like), oligodendrocyte progenitor-like (OPC-like), and astrocyte-like (AC-like) states within the tumor, respectively. In contrast, the tumor microenvironment (TME) primarily influences the MES-like state, indicating its distinct molecular characteristics and potential involvement in tumor-stroma interactions [4, 5].

The microenvironment surrounding cancer cells within the tumor, composed of vascular cells, immune cells, and non-cellular components like the extracellular matrix (ECM), is known as the TME. In recent years, the cellular and molecular composition of the TME has gained significant importance in the context of various anti-tumor therapies, including immunotherapy, across different types of cancer. The glioma TME is characterized by high levels of immunosuppression, setting it apart from the TME of other cancers [6]. While immunotherapy strategies, including immune-checkpoint inhibitors (ICIs), have shown promise in improving the prognosis of many cancers [7], they have not yielded substantial clinical benefits for GBM patients, primarily due to the immunosuppressive characteristics of the GBM TME [8]. The bidirectional interactions between cancer cells and the diverse cell populations present in the TME, facilitated by ligand-receptor (L-R) interactions, regulate multiple aspects of tumor behavior, ranging from immune suppression to angiogenesis and metastasis control [4, 9]. Among immune cells, tumor-associated macrophages (TAMs), including bone marrow-derived macrophages (BMD-TAMs) and microglia (MG-TAMs), have been recognized as the primary factor in the glioma TME [10]. A previous study has demonstrated that tumors with high levels of TAMs are more likely to develop MES-like states. Additionally, *NF1* mutations promote TAM recruitment and create a TAM-rich TME [11]. However, our understanding of the TME-derived signals and the mechanisms underlying the communications between tumors and TAMs in different categories of glioma remains incomplete.

In this study, we integrated single-cell RNA-seq (scRNA-seq) and bulk transcriptomes to dissect TAM heterogeneity across IDH-stratified gliomas and to identify TAM states that contribute to worse prognosis and ICI failure. We combined scRNA-seq with spatial transcriptomics (ST) and derived a bidirectional tumor-TAM L-R interactive network. Our work provides insights into the origins of GBM immunosuppressive and metastatic phenotypes.

## METHODS

### Preprocessing of scRNA-seq data

Preprocessing was performed using the Scanpy package v.1.39 [12]. Prior to analysis, the scRNA-seq dataset underwent quality control and filtering. Genes expressed in fewer than three cells were removed. Additionally, cells with less than 200 expressed genes and those with a mitochondrial content exceeding 25% were excluded. Subsequently, the data was normalized using a scaling factor of 10,000 followed by log transformation. Principal component analysis (PCA) was performed using the sc.pp.pca function and ‘n_comps’=50. Cells from patient number 6 were completely excluded due to low quality. To perform dimensionality reduction and batch correction, we employed the scVI model implemented in scvi-tools v.16.4 [13, 14]. We trained the scVI model on the raw counts of the 8000 highly variable genes (HVGs) with the following parameters: batch_key=‘sampleid’, n_latent=20, gene_likelihood=‘nb’, use_layer_norm=‘both’, use_batch_norm=None, encode_covariates=True, dropout_rate=0.2, n_layers=2. The training process ran for 300 epochs. Subsequently, a k-nearest neighbor (KNN) graph was constructed based on the similarity in the scVI latent space, using a k value of 50. Cells were clustered using the Leiden algorithm (scanpy.tl.leiden) with resolution = 1.5. Estimation of somatic copy number alterations (SCNA) from scRNA-seq data was performed using InferCNV v.1.6.0 (https://github.com/broadinstitute/inferCNV). Malignant cell states were classified using signatures as described in the original studies [15, 16] (Table S1) and using the function ‘AddModuleScore’ from Seurat R package v.4.3.0 [17]. Single-cell activity for MSigDB-hallmark (https://www.gsea-msigdb.org/gsea/msigdb/index.jsp) and Neftel et al. gene sets were determined using AUCell v.1.12.0 [18] package in R.

### Differential expression and gene set enrichment analysis

Differential expression analyses between BMD-TAM and MG-TAM clusters of IDH-WT and IDH-Mut conditions were performed using the diffxpy python package v.0.6.2 (https://diffxpy.readthedocs.io/en/latest/index.html). Differential expression was conducted via a two-sided Wald test that fits a negative binomial model to the raw count data using *IDH_status* covariate per gene for all genes expressed in at least 30 cells in the cluster of interest. The Benjamini-Hochberg method was performed for multiple testing correction. Overrepresentation analysis for MSigDb Hallmark gene sets (v.2020) was performed using the EnirchR interface implemented in GSEApy (https://github.com/zqfang/GSEApy). Associated p-values were computed with Fisher’s exact test. Adjusted p-values (q-values) were calculated using the Benjamini–Hochberg method for correction for multiple hypothesis testing.

### SCISSOR analysis

We utilized SCISSOR R package v.2.1.0 [19] to integrate phenotypic data of bulk RNA-seq datasets with scRNA-seq data. We downloaded gene expression data, along with corresponding survival and mutation data, for GBM and lower grade glioma (LGG) cohorts from The Cancer Genome Atlas (TCGA) on the GDC portal. SCISSOR was implemented in accordance with the guidelines provided in the accompanying vignettes. Our dependent variables were derived from two sources: genotypic data, which was analyzed using logistic regression, and patient overall survival rates, which were examined through Cox regression analysis. Based on the signs of the estimated regression coefficients, the cells with positive and negative association with the phenotype were indicated as Scissor^+^ and Scissor^-^ respectively. Cells with coefficients of zero were indicated as background cells. To validate the identified results, reliability significance test was performed. For each cell-type, we computed the log2-ratio of Scissor+ and Scissor^-^ cells as the mean fraction of Scissor^+^ cells vs. the mean fraction of Scissor^-^ cells.

### SCENIC analysis

We employed the single cell regulatory network inference and clustering reimplemented in python (pySCENIC) v.0.12.1 [20] package to unravel the gene regulatory networks (GRNs) of the myeloid cell populations within scRNA-seq dataset. As input, we provided the raw gene expression matrix and utilized the Lambert 2018 transcription factor (TF) list as a reference. The Grnboost2 algorithm within pySCENIC then constructed an adjacency matrix based on co-expression scores. In the next step, cisTarget was used to identify enriched motifs and infer regulons. We used the hg38_refseq-r80 500bp_up_and_100bp_down_tss and hg38 refseq-r80 10kb_up_and_down_tss motif databases using default settings for cisTarget. The cellular enrichment of each regulon was estimated by their area under the curve (AUC) scores calculated by AUCell. To visualize the results, AUC scores were added to the AnnData object and were plotted by the uniform manifold approximation and projection (UMAP).

### scRNA-seq pseudotime calculation and trajectory inference

We used Palantir v.1.3.1 [21] to generate pseudotime trajectories. We chose an early cell for trajectory (ID AACCTTTAGGCTTTCA-1-4) located at the maximal point of monocyte cluster in the UMAP. The terminal cells were automatically identified by the software. Default parameters were used.

CellRank v.1.5.1 [22] was employed as a downstream analysis for single-cell fate mapping and trajectory inference. Pseudotimekernel was used to compute cell-cell transition based on the precomputed Palantir trajectory and the result was projected onto the UMAP with arrows displaying RNA-velocity-like directionality towards the end of the pseudotime.

### Cell-cell interaction analysis

We utilized CellPhoneDB (https://github.com/Teichlab/cellphonedb) which is a curated database of ligands and their receptors with integrated statistical tools, to identify L-R interactions in scRNA-seq data from IDH-WT (n=5) and IDH-Mut (n=5) human glioma samples. Specifically, we employed Method 3 from CellPhoneDB v.3, which relies on the analysis of differentially expressed genes (DEGs) between cell-types.

### Bulk RNA-seq preprocessing and analysis

Raw FASTQ files underwent quality assessment with FastQC v.0.12.1 (https://github.com/s-andrews/FastQC). Adapters and low-quality bases were trimmed from the reads using TrimGalore v.0.6.10 (https://github.com/FelixKrueger/TrimGalore) with default parameters. Trimmed FASTQ files were then aligned to the human reference genome (hg38) with Salmon [23] for transcript quantification. Then, a gene-by-sample count matrix was generated from the aligned reads and was subjected to variance-stabilizing transformation (VST). PCA was performed on the transformed counts, and PCA plots were visualized using the top 1,000 most variable genes across samples. To correct for batch effects by donor, the removeBatchEffect function in the limma package (RRID:SCR_010943, v.3.36.5) was applied. Differential analysis of gene expression was performed using the DESeq2 (RRID:SCR_015687, v3.18) package in R.

### Signature scoring in bulk RNA-seq samples

The single sample gene set enrichment (ssGSEA) scores in bulk datasets were generated with the GSVA R package v.1.50.5 [24] using the ‘ssgsea’ method and ssgsea.nrom = T.

### Survival analysis

Survival analysis was conducted using the Cox proportional hazards (CoxPH) regression model implemented in the R package survival v.3.6. To evaluate the association between signature scores and survival outcomes, Kaplan-Meyer curves were generated using the survminer v.0.4.9 package in R. The Kaplan-Meyer curves depicted the survival probability for the top 25% and bottom 25% of samples stratified by their signature score.

### Analysis of ST data

We processed Visium ST data using Scanpy in Python, following the provided vignettes. Basic filtering of spots was performed based on total count and number of expressed genes. The data was then normalized using a scaling factor of 10,000 followed by log transformation. Variable gene detection was implemented using the ‘Seurat’ flavor. Dimensionality reduction, neighborhood graph construction, and UMAP calculation were subsequently performed. Each ST slide was analyzed independently.

To spatially map annotated cell types in scRNA-seq data onto the Visium ST datasets, we employed cell2location v.0.1.3 [25]. This software utilizes a Bayesian model to decompose ST data into spatially resolved estimates of cell type abundance. cell2location first employs negative binomial regression to estimate cell-type signatures from the reference scRNA-seq profiles. Subsequently, it leverages these derived signatures to perform non-negative decomposition of mRNA counts measured at each spatial spot.

Spatial cell-cell interaction analysis was performed using stLearn v.0.4.12 [26] in python. We leveraged the pre-curated L-R gene pairs included within the connectomeDB (v.2020) accessible through stLearn. The st.tl.cci.run function was employed to identify spatial spots exhibiting statistically significant co-expression of L-R gene pairs. This analysis utilized a permutation test with a pre-defined significance threshold of p-value < 0.05 and employed the default parameters within stLearn. Finally, the cell-cell interaction score for each significant L-R pair was determined using the lr_scores function.

### Human subjects and ethical considerations

Fresh glioma samples were obtained from surgical specimens undergoing resection at the Neurosurgery ward of Shariati hospital affiliated with Tehran University of Medical Sciences (TUMS). All patients provided written informed consent, in compliance with the institutional guidelines set by TUMS, prior to the collection of samples. Demographic details are provided in Table S2. Methodologies were executed in strict adherence to the principles outlined in the Declaration of Helsinki. Ethical clearance for this investigation was granted by the Institutional Review Board of TUMS, as confirmed by the ethical approval codes: IR.TUMS.SHARIATI.REC.1402.112 and IR.TUMS.SHARIATI.REC.1402.113.

### RNA extraction, cDNA synthesis and RT-qPCR

Total RNA was extracted from frozen tissue samples using TRIzol Reagent (Bio Basic Inc., Markham, Ontario, Canada) following the manufacturer’s protocol. RNA quality and purity were assessed using agarose gel electrophoresis and Nanodrop 2000 spectrophotometer (Thermo Scientific). cDNA was synthesized from the isolated RNA using the PrimeScript RT reagent (Takara Bio Inc., Shiga, Japan) according to the manufacturer’s protocol. RT-qPCR was performed using the AMPLIQON 2x qPCR Master Mix Green-No ROX using a LightCycler^®^ 96 System (Roche Life Science, Germany) according to the manufacturer’s instructions. Gene-specific primers used for amplification are listed in Table S3. Following RT-qPCR, gene expression levels were calculated using the 2^−ΔΔCt^ method with normalization to *TBP* and *B2M* as reference genes.

## RESULTS

### scRNA-seq reveals the heterogenic landscape of myeloid compartment in glioma TME

We used a publicly available dataset of 55,284 single cell transcriptomes from IDH-WT (n=5) and IDH-Mut (n=6) glioma samples. Following quality control and filtering down to 38,150 cells (Fig. S1A), we identified 9 major clusters using the UMAP dimensionality reduction technique (Fig. 1A left). Tumor and normal cells were segregated based on the inferred SCNA (Fig. S1B). Employing the Neftel-Suvà nomenclature [16], we classified malignant cells from IDH-WT samples into different stem-like states, including NPC-like and OPC-like, along with more differentiated states resembling AC-like and MES-like programs (Fig. S1C). We also adopted Venteicher-Suvà defined programs [27] to label IDH-Mut malignant cells into AC-like, oligodendrocyte-like (OC-like) and Stem-like states (Fig. S1D). Among non-immune (*CD45*^*-*^) cell types, we observed oligodendrocytes (*MOG*^*+*^), endothelial cells (CLDN5^+^), fibroblasts and pericytes (*DCN*^*+*^). The CD45^+^ immune cell populations comprised T-cells (*CD3D*^*+*^), B-cells (*CD79A*^*+*^), natural killer (NK) cells (*NKG7*^*+*^), and myeloid cells (*CD14*^*+*^, *FCGR3A*^*+*^) (Fig. 1A left; Fig. S1E).

**Fig 1.**
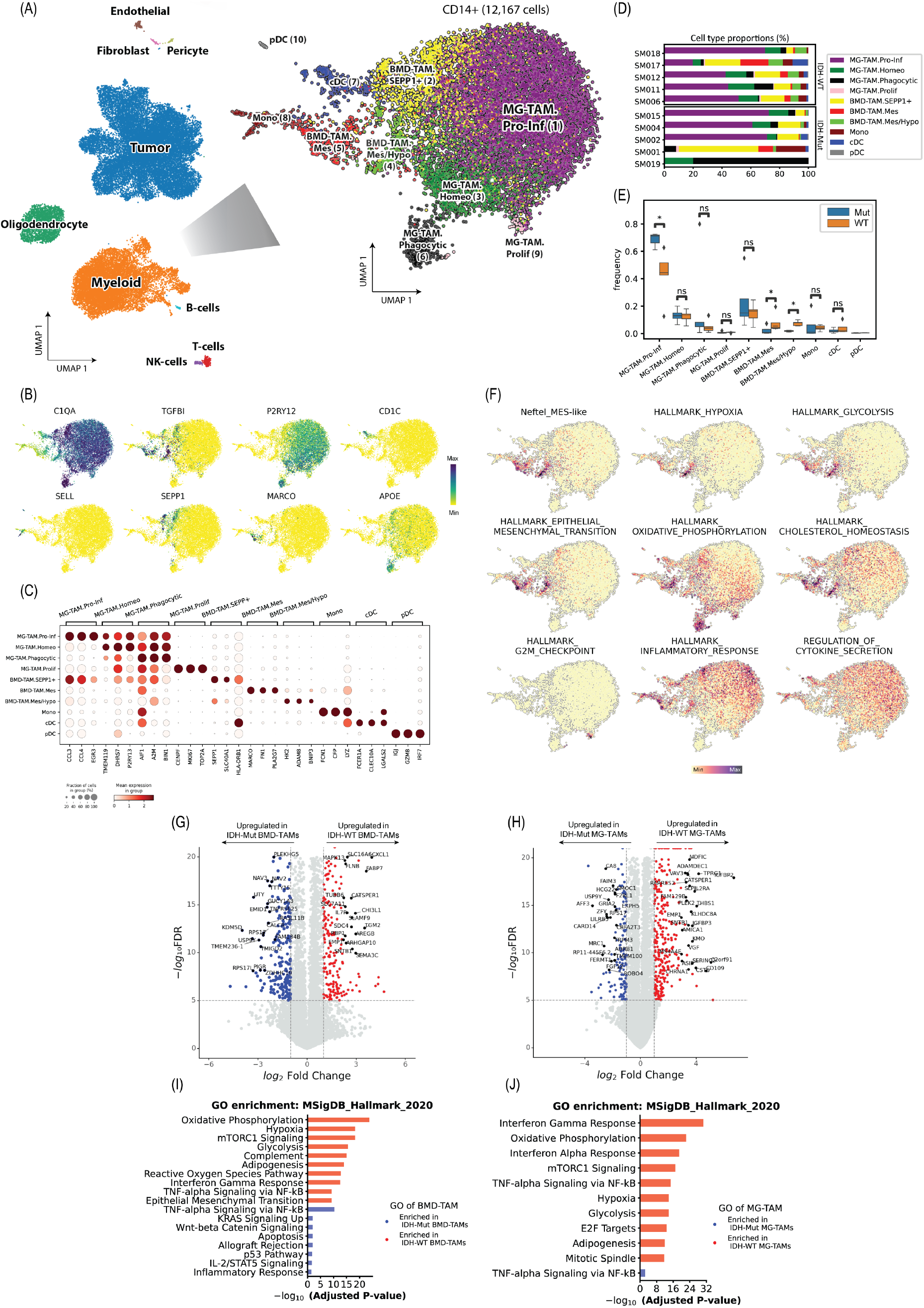
Dissecting functional states of TAMs within glioma TME using scRNA-seq. (A) UMAP plots of whole tumor (left) and myeloid compartment (right) colored and labeled by subcluster annotations; (B) UMAP plots showing expression levels of marker genes in myeloid compartment; (C) Bubble heatmap illustrating the expression of marker genes in the represented myeloid subsets. The dot size indicates the fraction of expressing cells and are color coded based on the average expression levels; (D) Bar plots demonstrating the proportion (%) of myeloid cell states assigned to the glioma patients; (E) Box plot showing the relative frequency of myeloid subtypes in IDH-WT and IDH-Mut samples. P-values were calculated unpaired Wilcoxon rank-sum test; (F)UMAP plots depicting the scaled AUC scores, highlighting cellular activity levels across hallmark gene sets; (G, H) Volcano plots showing DEGs between (G)BMD-TAMs and (H) MG-TAMs of IDH-WT and IDH-Mut tumors. The statistical p-values were determined by the two-tailed Wald test with Benjamini and Haschberg correction for multiple testing. Selected represented genes are highlighted. The color coding represents an adjusted p-value below 0.01 and |log2 fold change| above 1.5; (I, J) Overrepresentation analysis for MSigDB_Hallmark_2020 gene sets using up and down regulated genes in (I) BMD-TAMs and (J) MG-TAMs of IDH-WT, IDH-Mut tumors. p-values were determined using Fisher’s exact test.

To effectively decipher functional heterogeneity of the *CD14*^*+*^ myeloid cluster with 12,167 cells, we first evaluated the expression of canonical biomarkers and then measured the activity of transcriptomic programs via AUCell software [18] (Table S4, S5). As a universal biomarker of mature macrophages, *C1QA* separated TAMs from non-TAMs (Fig. 1B). Cluster 8 exhibited significant upregulation of marker genes characteristic of classical monocytes, including *SELL, FCN1* and *VCAN* [28] (Fig. 1B, C). Cells expressing microglial signature genes (e.g., *TMEM119, P2RY12, CX3CR1*) were identified as MG-TAM (Fig. 1B). In contrast, *TGFBI and CLEC12A* marker genes were associated with BMD-TAMs (Fig. 1B). We identified two subpopulations of BMD-TAMs marked by increased activity for the MES-like gene set (Fig. 1F) [16]. These cells appear to correspond to the recently described mesenchymal myeloid phenotype [4]. Interestingly, one subcluster demonstrated higher enrichment for hypoxia and glycolysis programs, while the other displayed increased activity for oxidative phosphorylation and tumor-supportive genes including *FN1* and *MARCO* (Fig. 1C). We annotated these clusters as BMD-TAM.Mes/Hypo and BMD-TAM.Mes respectively. EMT and cholesterol homeostasis modules were enriched in both clusters (Fig. 1F). Cluster 2 contained a subpopulation of BMD-TAMs expressing high levels of *SEPP1, FOLR2*, and *SLC40A1* – genes often linked to anti-inflammatory activation [28] (Fig. 1B, C; Fig. S1H). We designated these cells as BMD-TAM.SEPP1^+^ (Fig. 1A right).

We explained the functional heterogeneity of MG-TAMs by four programs: An MG-TAM subpopulation exhibited elevated expression of homeostatic microglia markers (e.g., *TMEM119, P2RY13*) and concomitantly displayed lower expression of genes associated with catabolic processes (e.g., *GPX1*) or cell activation (e.g., *CCL3L3*) (Fig. 1C; Fig. S1F), suggesting a homeostatic low-activation state. Therefore, we annotated these cells as MG-TAM.Homeo. A small fraction of MG-TAMs in cluster 9 exhibited elevated expression of G2M and S phase genes (Fig. 1F), such as *MKI67* and *STMN1*, signifying a proliferative phenotype. According to Yeo *et al*. Ki67+ proliferating MG-TAMs increase only in tumoral microglia upon GBM progression and adopt a different transcriptome compared to non-proliferating MG-TAMs [29]. MG-TAMs in Cluster 1 displayed an M1-like pro-inflammatory phenotype, characterized by increased expression of *EGR3* and *CCL3* genes (Fig. 1C). This cluster demonstrated enrichment for hallmark gene sets of proinflammatory response and cytokine secretion (Fig. 1F). Cluster 6 exhibited an upregulation of genes (e.g. *APOE, SPP1, BIN1, LGALS3, FABP5, GPNM, AIF1, CD68*) and also opsonins (e.g. *C1QA, C1QB, C1QC*) linked to phagocytic and lipid-associated macrophages [30-32] (Fig. 1 B, C; Fig. S1G). These genes are also similar to markers of aging microglia [33].

We compared the cellular composition of myeloid compartment between IDH-WT and IDH-Mut samples. The MG-TAM.Pro-Inf cluster emerged as the largest in both groups, with a higher frequency observed in IDH-Mut tumors (Fig. 1E). Conversely, The BMD-TAM.Mes cluster was predominantly found in IDH-WT samples although one IDH-Mut patient exhibited a high percentage as well (Fig. 1D, E). Similarly, the proportion of BMD-TAM.Mes/Hypo cells was also significantly higher in IDH-WT patients. No other clusters demonstrated significant compositional differences (Fig. 1E).

To further investigate the differences in TAM populations between IDH-WT and IDH-Mut gliomas, we conducted a pseudobulk differential expression analysis. This analysis identified DEGs in both conditions per BMD-TAM and MG-TAM clusters (Fig. 1G, H; Table S6, S7), supporting the existence of distinct cell states. Gene ontology enrichment of up-regulated genes in IDH-WT BMD-TAMs revealed enrichment of pathways related to oxidative phosphorylation, hypoxia as well as mTORC1 signaling (Fig. 1I). IDH-WT MG-TAMs were found to be involved in Interferon Gamma and Alpha response, and oxidative phosphorylation (Fig. 1J). We collectively highlight that TAMs in the TME of IDH-WT GBM, exhibit enhanced interferon activity and a heightened response to hypoxia.

### Mesenchymal BMD-TAMs are negatively associated with patient survival and immunotherapy response in GBM

Next, we set out to complement our scRNA-seq analysis by integrating genomic and survival information provided by the TCGA reference project. The relationship between genomic profile of the tumor and its TME contexture is intricate and complex. Research has shown that mutations in certain genes can influence the GBM TME by altering expression of the factors that recruit, activate or suppress various immune and stromal cells in the TME. Previous investigations have integrated profiling techniques with functional studies on *PTEN*-deficient GBM models, elucidating that *PTEN* mutations promote BMD-TAM infiltration with no effect on MG-TAMs [34]. Alterations in *NF1* as a negative regulator of RAS/MAPK pathway drives mesenchymal transformation of GBM through modulation of AP-1 transcription factor *FOSL1* expression [35]. *NF1* deficiency is shown to trigger chemoattraction of BMD, MG-TAMs in GBM [11]. As the signature receptor of the classical subtype, *EGFR* is frequently amplified in GBM [36]. Full-length EGFR, alongside the truncated EGFRvIII, cooperates with KRAS to elevate expression levels of the chemokine CCL2, thereby facilitating macrophage infiltration in GBM mouse models [37]. *TP53* gain-of-function mutations in human GBM cell lines trigger activation of NF-κB signaling, leading to increased expression of *CCL2* and *TNFα*. These chemokines, in turn, recruit BMD, MG-TAMs into GBM TME [38]. Characterizing the link between tumor cell-intrinsic genetic events and the TAM composition could inform development of personalized immune intervention strategies in glioma.

Using the SCISSOR package in R, we evaluated association of transcriptomic signatures of myeloid components with genotypic information across GBM (n = 154) and LGG (n = 507) TCGA bulk RNA-seq datasets. According to our results, mutations in all four mesenchymal signature genes—*NF1, PTEN, RB1* and *TP53*—were associated with BMD-TAM.Mes cluster (Fig. 2A, B, C, E). However, only *NF1 and PTEN* were also associated with the BMD-TAM.Mes/Hypo cluster (Fig. 2A, B). Interestingly, alterations in any of these genes did not seem to have a noticeable effect on SEPP1^+^ BMD-TAM cluster. Relating to MG-TAM clusters, *TP53* mutations was positively associated with MG-TAM.Pro-Inf (Fig. 2E). *IDH1* mutations in the LGG cohort had also positive impact on this cluster (Fig. 2F). Patients with *PTEN* deficiency exhibited positive associations with MG-TAM. Phagocytic (Fig. 2B). *RB1* and *TP53* mutations demonstrated positive associations with MG-TAM.Prolif but negative associations with MG-TAM.Phagocytic (Fig. 2C, E). We also observed distinct infiltration patterns in the cDC cluster: patients with *PTEN, RB1*, and *EGFR* mutations demonstrated high infiltration, while those with *TP53* and *IDH1* mutations had low infiltration. A similar trend was observed in the monocyte and pDC clusters, where the *TP53* genotype was associated with high enrichment, and the *NF1* and *PTEN* genotypes were linked to low infiltration (Fig. 2A-F).

**Fig 2.**
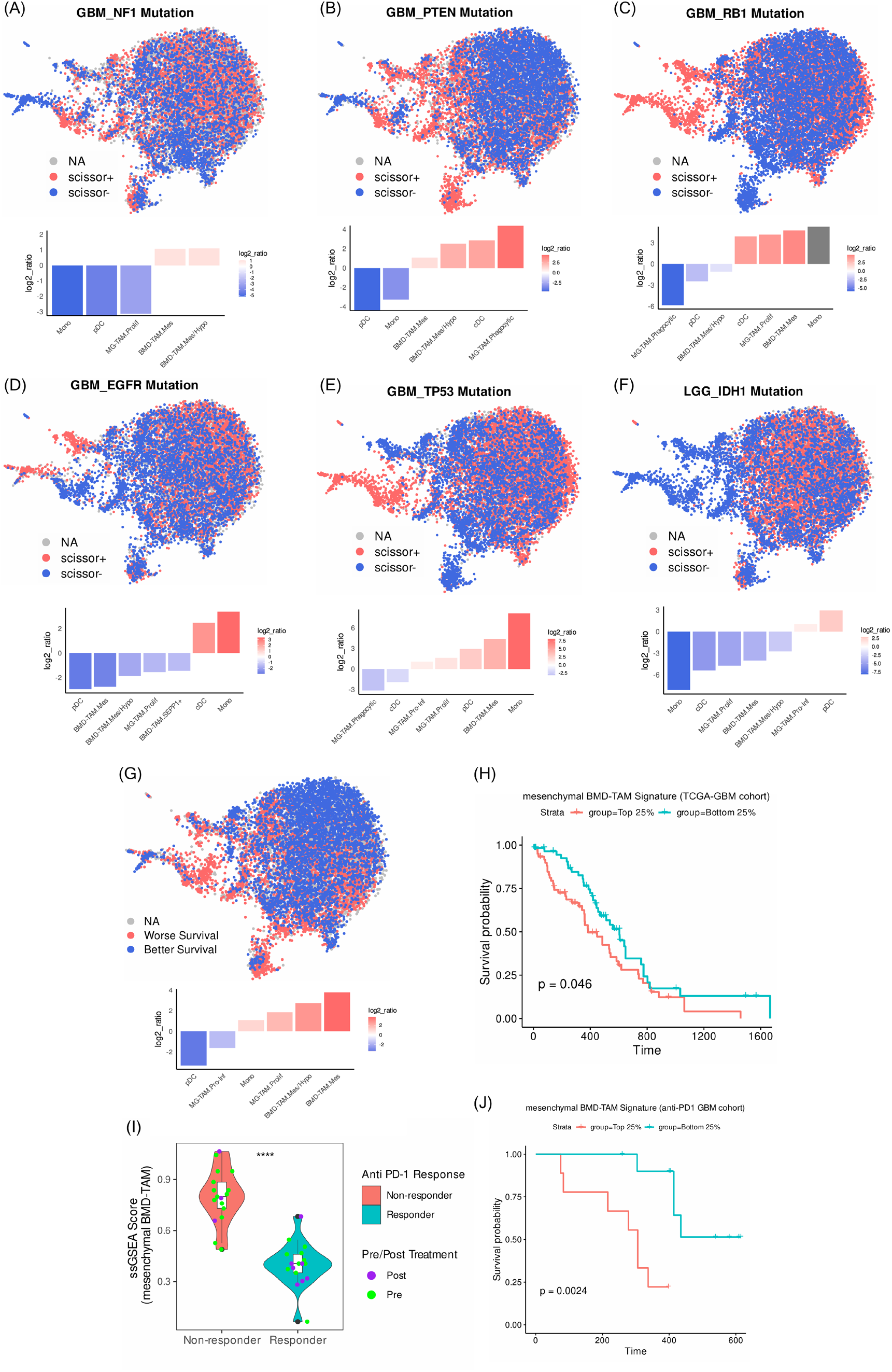
Glioma myeloid cell-type composition associates with patient genotype, overall survival and response to ICI therapy. (A-G) UMAP plots showing selected cells by SCISSOR analysis. Dots are colored according to the association of cells with the phenotype (blue: negative, red: positive). Shown are clusters with |log2 ratio| > 1; (A) Association of cellular composition of myeloid cluster with *NF1* mutations in GBM patients (n = 17); (B) Association of cellular composition of myeloid cluster with *PTEN* mutations in GBM patients (n = 53); (C) Association of cellular composition of myeloid cluster with *RB1* mutations in GBM patients (n = 11); (D) Association of cellular composition of myeloid cluster with *EGFR* mutations in GBM patients (n = 41); (E) Association of cellular composition of myeloid cluster with *TP53* mutations in GBM patients (n = 52); (F) Association of cellular composition of myeloid cluster with *IDH1* mutations in GBM patients (n = 394); (G) Association of cellular composition of myeloid cluster with patient overall survival; (H) Kaplan-Meier survival curve showing clinical relevance of mesenchymal BMD-TAM signature in patients with high (upper quartile) and low (lower quartile) enrichment scores in TCGA-GBM cohort. Statistical p-value was determined using CoxPH regression; (I) Bar plots with surrounding violins demonstrating the GSVA of DEGs using mesenchymal BMD-TAM signature in non-responder vs. responder patients treated with nivolumab or pembrolizumab. Center line represents mean score. P-value was calculated using two-tailed Wilcoxon rank-sum test; (J) Kaplan-Meier plot illustrating the comparison of overall survival rates in GBM patients undergoing treatment with either nivolumab or pembrolizumab, categorized by high (upper quartile) and low (lower quartile) enrichment scores for the mesenchymal BMD-TAM signature.

We proceeded to evaluate association of transcriptomic signatures of cell types with survival outcomes in 236 patients from the TCGA-GBM cohort. Patients with tumors enriched for BMD-TAM.Mes and BMD-TAM.Mes/Hypo clusters showed a significant association with poorer survival, whereas MG-TAM.Pro-Inf and pDCs were the strongest positive survival indicators. BMD-TAM.SEPP1^+^ and MG-TAM.Phagocytic clusters seemed to have no significant impact on overall patient survival (Fig. 2G). Subsequently, we curated a signature gene set (Table S8) as a union of genes with highly specific expression within two subclusters of BMD-TAM.Mes and BMD-TAM.Mes/Hypo and then investigated its connection with patient survival. Notable, Kaplan-Meier survival analysis demonstrated an apparent association between enrichment of mesenchymal BMD-TAM gene set and unfavorable overall survival of patients from TCGA-GBM (Fig. 2H).

To further evaluate the relevance of mesenchymal BMD-TAMs in clinical settings, we used a longitudinal bulk RNA-seq dataset of GBM patients (n=17) treated with PD-1 checkpoint inhibitors (nivolumab or pembrolizumab). The original study showed that the non-responder patients were enriched for *PTEN* mutations, suggesting that these mutations may induce a distinct immunosuppressive TME correlated with therapy failure [39]. We found mesenchymal BMD-TAM signature highly enriched in non-responder patients compared to responders (Fig. 2I), indicating a potential association of these cells with anti-PD1 therapy failure. Furthermore, survival analysis within this cohort demonstrated that patients with high enrichment of the mesenchymal BMD-TAM signature exhibited significantly worse overall survival (Fig. 2J). To address the urgent need for therapeutic targets against these TAM clusters, we propose SLAM family member 9 (SLAMF9) as a promising candidate. SLAMF9 is a member of the derived gene signature associated with mesenchymal BMD-TAMs. It encodes a cell surface receptor specifically expressed on mesenchymal BMD-TAMs, while being absent in healthy central nervous system (CNS) tissues (www.proteinatlas.org) (Fig. S2B, C, D). SLAMF9 is involved in the production of proinflammatory cytokines and migratory patterns in both murine and human melanoma TAMs [40].

### Inference of gene regulatory networks predicts TFs governing myeloid cells

Next, we applied pySCENIC [18, 20] to comprehensively reconstruct GRNs for all myeloid cells in the GBM TME. This approach can potentially enhance our knowledge of the functions and regulatory mechanisms governing different myeloid subsets, which could lead to new clinical implications and therapeutic strategies. By linking *cis-regulatory* sequence information with single cell transcriptomes, pySCENIC deciphers critical TFs and their target genes, which together form a regulon. Our analysis on scRNA-seq of IDH-WT GBM identified regulons that define the GRNs of the myeloid compartment. To rank these regulons by their cell-type specificity, we employed Regulon Specificity Score (RSS) plots, a metric based on Jensen-Shannon divergence.

MG-TAM.Pro-Inf were governed by two zinc finger TFs EGR2, EGR3 and several immediate early response TFs including FOS, JUN and JUNB (Fig. 3A). MG-TAM subsets with elevated levels of chemokine genes (e.g. *CCL2, CCL4*) and early growth factor 2 and 3 (*EGR2, EGR3*) have been denoted as ‘pre-activated’ [41]. In a recent study on inflammaging, Karakaslar *et al*. demonstrated that the age-associated elevated expression of AP-1 members of TFs, particularly Fos, Jun, and Junb, was conserved across mice immune tissues including macrophages, potentially contributing to the increased inflammation observed with aging [42]. Five top regulons of MG-TAM.Phagocytic were SPI1, GABPA, POLE4, ETS1 and USF2. The SPI1 (also known as PU.1) serves as a central regulator of myeloid cells, governing both the development and function of microglia. Microglia with reduced *Spi1* expression exhibit diminished phagocytic capacity [43], while conversely, increased *Spi1* levels enhance their ability to engulf zymosan particles [44]. MG-TAM.Prolif displayed high activity for POLE3, TFDP1, MAZ, RAD21, and NFIA regulons. The *TFDP1* gene encodes the heterodimeric partner DP1 of the E2F, which is crucial for cell proliferation as it activates a set of growth-related genes [45] (Fig. 3A).

**Fig 3.**
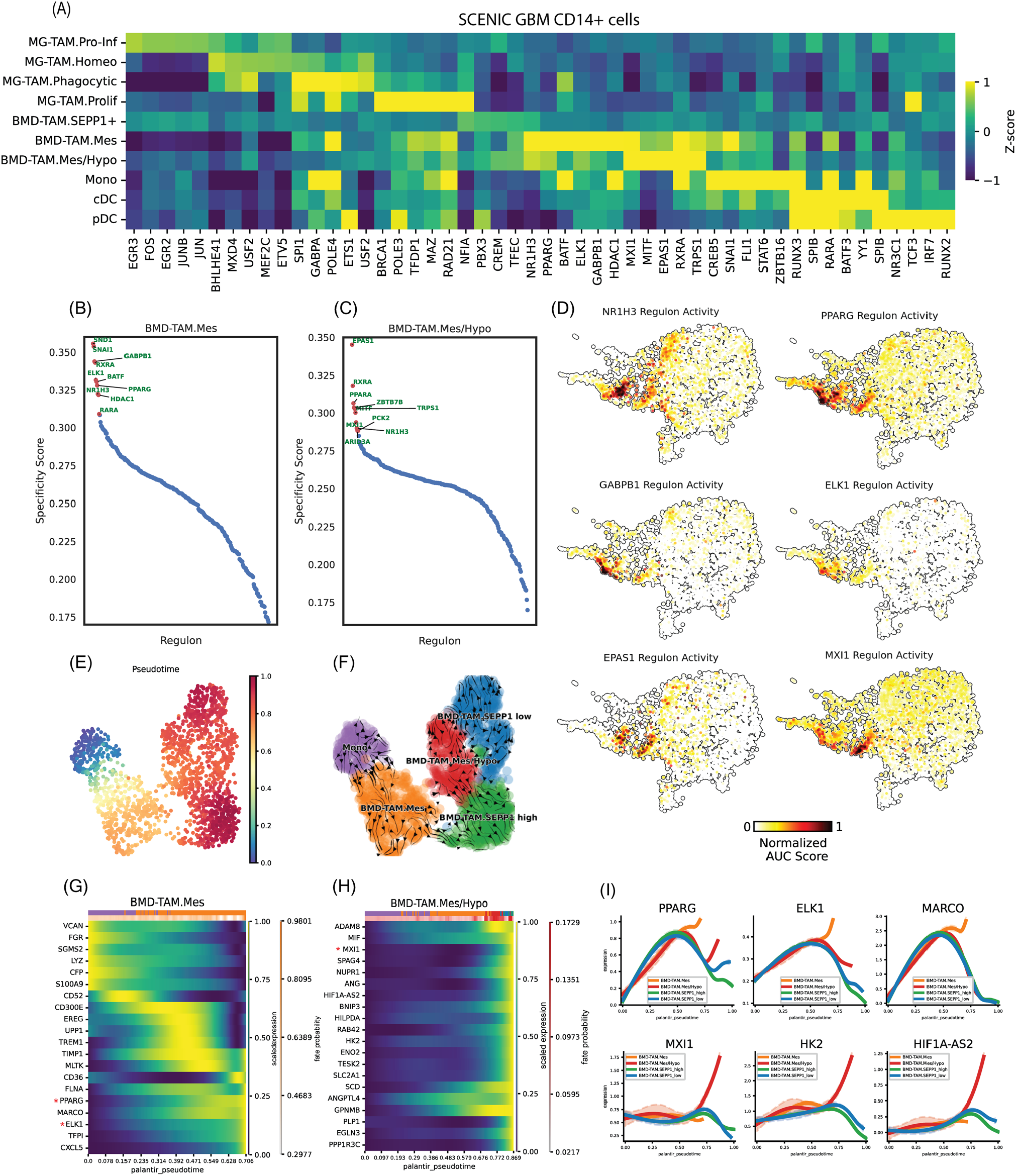
GRN inference and trajectory analysis. (A) Enriched TF activity across myeloid cells in IDH-WT GBM, identified by the pySCENIC algorithm, and depicted as a heatmap of cell-type median relative z-scores; (B, C) RSS plots for pySCENIC-derived regulons in (B) BMD-TAM.Mes and (C) BMD-TAM.Mes/Hypo clusters; (D) UMAP plots illustrating the normalized AUC scores of NR1H3, PPARG, GABPB1, ELK1, MXI1 and EPAS1 regulons; (E) UMAP visualization of pseudotime generated by Palantir; (F) Streamlines visualizing the direction of cell-cell transition calculated based on pseudotime; (G, H) Heatmap plots demonstrating the inferred top-ranked driver genes highly correlated with fate probabilities of (G) BMD-TAM.Mes and (H) BMD-TAM.Mes/Hypo clusters; (I) Smoothed expression trends of representative genes along the pseudotime. The trend for each gene is shown for each trajectory leading to the indicated terminal population.

Among the top regulons inferred active in BMD-TAM.Mes were PPARG (also known as NR1C3) while PPARA was specific to BMD-TAM.Mes/Hypo. RXRA was active in both BMD-TAM.Mes and BMD-TAM.Mes/Hypo (Fig. 3A, B, C). By forming heterodimers with retinoid X receptor α (RXRα), PPARγ contributes to the polarization of macrophages into an immune tolerant state in which gluconeogenesis is dominant [46, 47] while PPARα modulates an immune reactive glycolytic state [48, 49]. Monocytes and BMD-TAM.Mes showed relatively high activity for BATF and SNAI1. NR1H3 (also known as LXRA) exhibited considerable activity across all BMD-TAM subsets. Activation of NR1H3 (LXRα) in human macrophages potentiates HIF-1α signaling and glycolysis [50], establishing a positive feedback loop that promotes foam cell formation [51]. LXRα also acts as a key modulator linking cholesterol accumulation in macrophages with suppression of inflammatory responses [52]. ELK1 regulon showed high activity and specificity in BMD-TAM.Mes cells. Tumor-derived lactate in colorectal cancer (CRC) induces expression of *ELK1*, promoting expression of the inhibitory immune checkpoint receptor Sirpα in TAMs and evasion of CRC from innate immune surveillance [53]. BMD-TAM.Mes/Hypo demonstrated the highest activity for the canonical hypoxia inducible factor EPAS1 (also known as HIF2A) and MXI1, a well-documented hypoxia-induced TF that is among the genes targeted by HIF1A [54]. Finally, we ranked the inferred regulons based on their RSS, highlighting the top ten most specific regulons. Accordingly, AUC scores for six transcription factors of NR1H3, PPARG, GABPB1, ELK1, MXI1 and EPAS1 with highest activity and specificity in mesenchymal BMD-TAMs were projected on UMAP (Fig. 3D).

Then, we sought to explore differentiation trajectories within the heterogenous population of BMD-TAMs. Using a cell in the monocyte cluster as the starting state, we employed the Palantir algorithm [21] for trajectory analysis and ordered BMD-TAMs along a high-resolution pseudotime scale, which reflects their progression through various developmental states. The pseudotime gradient, initiated with monocytes, advanced through BMD-TAM.Mes and BMD-TAM.Mes/Hypo as intermediate states and culminated in SEPP1-high and SEPP1-low branches of BMD-TAMs (Fig. 3E). Subsequently, we utilized the Cellrank2 modular framework to delve into the molecular dynamics of BMD-TAMs. By integrating other sources of input data such as cell-cell transcriptomic similarities, CellRank can address the noise in Pseudotime. We computed the cell-cell transition matrix using the pseudotime kernel and illustrated directionality of the trajectory inference (Fig. 3F). CellRank successfully identified monocytes as the initial state. Estimation of fate probabilities showed that most cells were fate biased towards the SEPP1_high terminal state (Fig. S3A, B). In order to identify driver genes of the BMD-TAM.Mes, BMD-TAM.Mes/Hypo, expression trends of genes whose relative expression showed the strongest correlation with terminal fate probabilities, were plotted against the pseudotime. Noteworthy among the top 20 driver genes were pySCENIC-derived TFs *PPARG* and *ELK1* in BMD-TAM.Mes and *MXI1* in BMD-TAM.Mes/Hypo (Fig. 3G, H). We overlaid the expression of master regulators and lineage associated markers to visualize trends based on calculated fate probabilities. Expression of *ELK1, PPARG*, and *MARCO* genes related to BMD-TAM.Mes and *MXI1, HK2*, and *HIF1A-AS2* associated with BMD-TAM.Mes/Hypo were upregulated while approaching their associated terminals at pseudotime units around 0.6 and 0.8, respectively (Fig. 3I).

### Analysis of cell-cell communication reveals GBM-specific tumor-TAM interactions involved in TAM recruitment, proliferation, immunosuppression and metastasis

Using CellPhoneDB L-R complexes repository, we investigated differences in intercellular crosstalk between tumor cell-states and TAMs among the two major categories IDH-WT and IDH-Mut gliomas (Fig. 4A). The spectrum of our inferred L-R interactome covered diverse IDH-WT-only signaling networks linked to immunosuppression, TAM recruitment, cellular differentiation and metastasis. The unique interaction observed in IDH-WT, mediated by the ANXA1 ligand expressed by the MES-like state towards FPR1 and FPR3 receptors, which were highly expressed on MG-TAMs and BMD-TAMs, respectively (Fig. S4A, B, G, H), has recently been reported to play an immunomodulatory role in GBM by recruiting and polarizing macrophages to suppress T-cells [55]. Furthermore, the VEGFA ligand released by MES-like cells significantly interacts with the NRP1 and NRP2 receptors highly expressed on MG-TAMs and BMD-TAMs (Fig. S4C, D, I, J). This interaction is believed to be crucial for TAM survival, recruitment, and establishment of an immunosuppressive TME [56]. According to Hara et al., macrophage-derived oncostamin M (OSM) ligand interacts with OSMR receptors on GBM cells, potentially inducing a MES-like phenotype by activating the STAT3 pathway [4]. FN1, which was highly expressed by BMD-TAM.Mes (Fig. S4E, K), forms various combinations with integrin receptors (ITGAV, ITGB1) on MES-like malignant states. A multitude of studies corroborate that such interactions foster tumor progression and metastasis [57]. Additionally, the TNFSF12 (also known as TWEAK) ligand secreted by MG-TAMs combines with the TNFRSF12A (also known as Fn14) receptor on MES-like and AC-like states, modulating glioma progression and proliferation by regulating the NF-κB pathway [58] (Fig. 5; Fig. S4F, L). In IDH-Mut, on the other hand, the CX3CL1-CX3CR1 axis mediated by the AC-like state and MG-TAMs or BMD-TAMs has been shown to negatively regulate glioma invasion likely via promoting tumor cell aggregation [59]. In addition, previous evidence has demonstrated that CSF1-CSF1R interaction, facilitated by AC-like state and BMD-TAMs or MG-TAMs, regulates the production, differentiation, and function of TAMs. While CSF1R inhibition has been proposed as an antitumor therapy in majority of cancers, its effectiveness in context of GBM has yielded inconclusive results [60]. In light of recent studies, APOE protein secreted by the AC-like cells binds to the TREM2 receptor on TAMs, particularly MG-TAMs, forming a combination that leads to immune suppression within the TME which correlates with tumorigenesis and T cell exhaustion [61, 62]. Moreover, VCAM1, situated on the surface of AC-like cells, has been demonstrated to interact with the integrin receptors of BMD-TAM, contributing to recruitment of TAMs along with promotion of pro-survival signals within tumor cells [63] (Fig. 4A).

**Fig 4.**
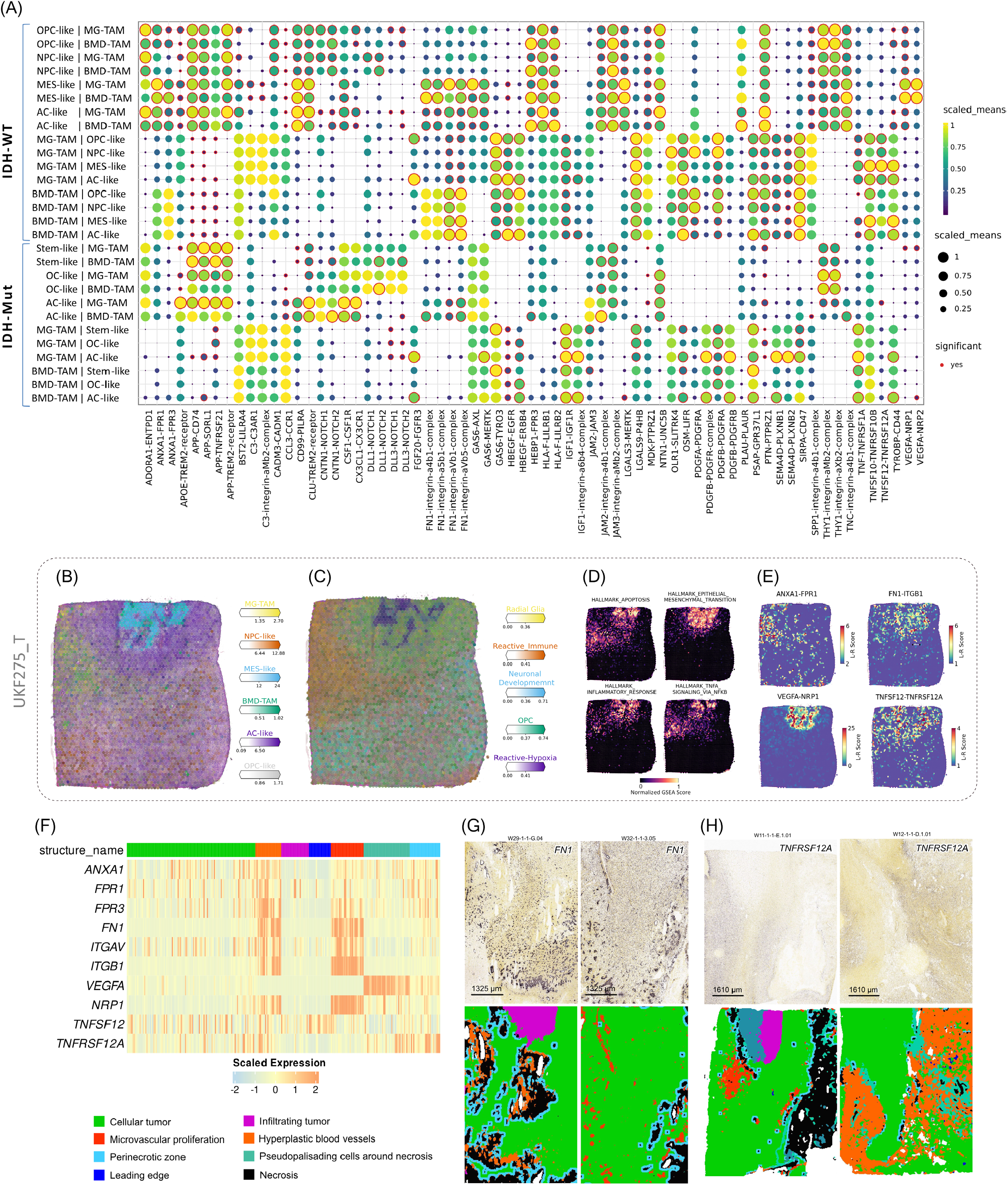
Inference of cell-cell communication between tumor cell states and TAMs. (A) Dot plot showing scaled mean expression of predicted ligand and receptors mediating interactions between tumor cell states and TAMs in IDH-WT and IDH-Mut tumors; (B) Surface plot showing the spatial locations of cell types in Visium ST datasets from UKF275_T donor; (C) Surface plot indicating spatial enrichment scores for five transcriptional programs; (D) Surface plot showing spatial GSEA scores for hallmark gene sets; (E) Spatial cell-cell interaction of representative L-R pairs; (F) Heatmap plot showing the scaled expression levels of *ANXA1, FN1, VEGFA, TNFSF12* genes along with their associated receptors in bulk RNA-seq datasets from seven anatomically distinct microregions provided by the IVY-GAP; (G, H) GBM RNA-ISH experiments provided by IVY-GAP; (G) RNA-ISH results showing expression of *FN1* gene in anatomically annotated regions; (H) RNA-ISH results showing expression of *TNFRSF12A* gene in anatomically annotated regions.

**Fig 5.**
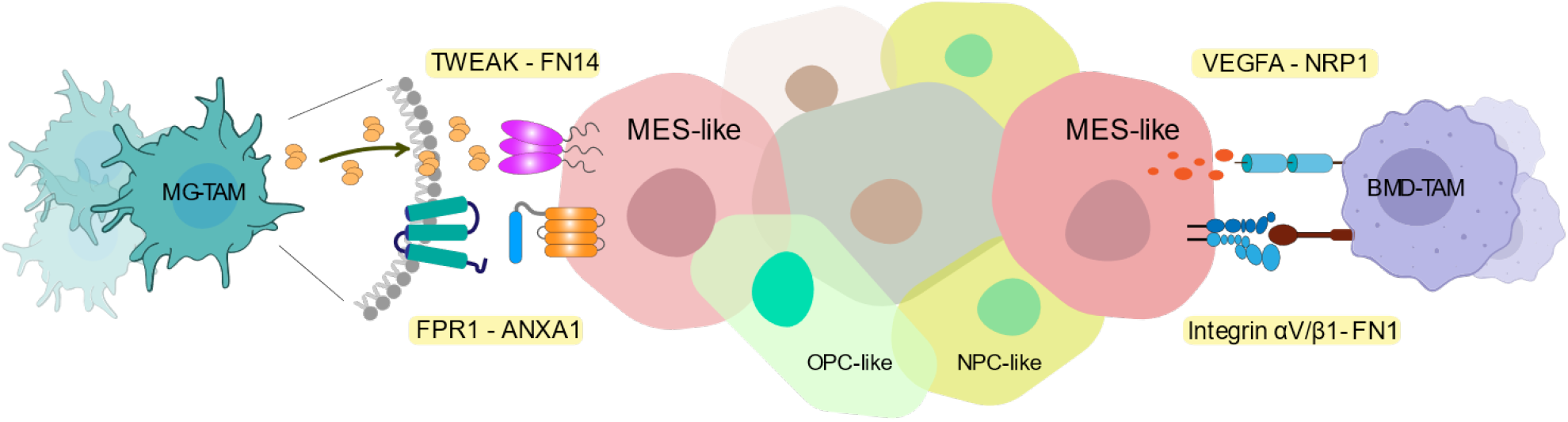
Schematic illustration of inferred L-R interactions between IDH-WT malignant states and TAMs.

To complement our scRNA-seq-derived findings, we set out to examine spatial organization of inferred communications using ST. Spatial omics techniques can provide valuable insights into the intricate architecture and cellular organization of the TME helping to further elucidate the spatial relationship of its components. Spatial cell-cell interaction analysis was conducted via stLearn [26]. In line with our scRNA-seq-derived results, ST analysis further indicated significant interaction of ANXA1 ligands with FPR1 receptors in reactive immune regions dominated largely by TAMs (Fig. 4B, C, E; Fig. S4M, N, P). Activation of FPR1 receptors through binding of the ANXA1 ligand has recently been demonstrated to shift the polarization of BMD, MG-TAMs towards a protumor M2 phenotype. These polarized immune cells then secrete cytokines like CCL22, recruiting T regulatory (Treg) cells, ultimately contributing to the formation of an immunosuppressive TME [64]. ST analysis confirmed significant FN1-ITGB1 and FN1-ITGAV interactions around the MES-like territory forming a distinct belt-like pattern (Fig. 4E; Fig. S4P). The reactive hypoxia area was featured with increased activity for the EMT-related gene set (Fig. 4D; Fig. S4O). Engagement of integrin heterodimers with FN1 molecules deposited in the ECM leads to subsequent receptor clustering and initiates accumulation of intricate adaptor and signaling protein clusters that regulate downstream integrin signaling pathways, typically by autophosphorylation of focal adhesion kinase (FAK) and subsequent recruitment of Src family kinase (SFK) [65]. FAK signaling in turn, governs various facets of the cancer cell such as cell adhesion, migration, protease expression, as well as survival and proliferation [66]. ST analyses demonstrated that NRP1-VEGFA interaction is predominant in the reactive hypoxia zone (Fig. 4C, E; Fig. S4N, P). VEGFA, a potent growth factor, promotes chemotaxis in TAMs by activating multiple signaling pathways, including the PI3K/Akt and MAPK cascades. Binding of VEGFA to its receptors, VEGFRs, and co-receptor NRP1 on the TAM surface triggers these intracellular signaling events [56, 67]. These pathways contribute to both TAM survival and regulation of cytoskeletal rearrangements, modulating the cellular machinery involved in cell movement. This coordinated response enables TAMs to migrate towards areas with increased vascularization or ongoing angiogenesis, where they can participate in processes that promote tumor growth and progression [68, 69]. Moreover, we observed significant interactions between the TNFSF12 ligands and the TNFRSF12A receptors within reactive-hypoxia regions (Fig. 4C, E; Fig. S4N, P). These areas exhibited an enrichment for hallmark gene signatures associated with TNFA signaling via NFKB and apoptosis (Fig. 4D; Fig. S4O). TWEAK-Fn14 signaling activates a variety of intracellular signaling pathways such as regulation of TNF-induced cell death and stimulation of classical and alternative NF-κB pathways [70, 71]. TWEAK-stimulated activation of the non-canonical NF-κB pathway and the NF-κB-inducing kinase upregulates *MMP-9* expression and promote invasion of GBM cells [72].

In follow-up, we used IVY-GAP anatomical transcriptional atlas [73] to evaluate expression of *ANXA1, VEGFA, FN1* and *TNFSF12* genes, along with their corresponding receptors, in 7 anatomical regions from GBM patients (n=10). Notably, microvascular proliferation (MVP) and hyperplastic blood vessel (HBV) regions related to angiogenesis, immune response regulation, and wound healing exhibited increased expression of *FN1, ITGAV* and *ITGB1* genes. *VEGFA* levels were elevated in pseudopalisading cells around necrosis (PAN), characterized by hypoxia and stress response. The *TNFRSF12A* gene was detected in most regions, with enrichment in the perinecrotic zone (PNZ). *TNFSF12* was highly expressed in leading edge (LE), which largely consists of non-neoplastic cells (Fig. 4F). RNA in-situ hybridization (RNA-ISH) data provided by the IVY-GAP corroborated these observations, demonstrating elevated expression of *FN1* in MVP and HBV and *TNFRSF12A* in PNZ (Fig. 4G, H).

### Bulk expression profiles reflect single-cell findings

To investigate whether the observed differences in cellular communications manifest at the scale of bulk tumor tissue, we employed bulk RNA-seq and compared mRNA levels of *ANXA1, FN1, NRP1* and *TNFRSF12A* genes between IDH-WT (n=162) and IDH-Mut (n=534) TCGA cases. Recapitulating the single-cell results, we found a negative association between *IDH1* mutation and expression of our candidate genes which were significantly upregulated in IDH-WT (Fig. 6A; Table S9). Importantly, tumors of mesenchymal (ME) transcriptomic subtype had the highest expression for all four genes, while the lowest expression was observed in the proneural (PN) GBM subtype (Fig. 6B). High expression levels of identified candidate genes indicated a negative prognosis for GBM patients (Fig. 6C-F). To corroborate these findings, we employed RT-qPCR on an independent cohort of IDH-WT (n=39) and IDH-Mut (n=41) patients. RT-qPCR analysis confirmed a significant upregulation of *ANXA1, FN1, NRP1* and *TNFRSF12A* genes in IDH-WT compared to IDH-Mut samples (Fig. 6G).

**Fig 6.**
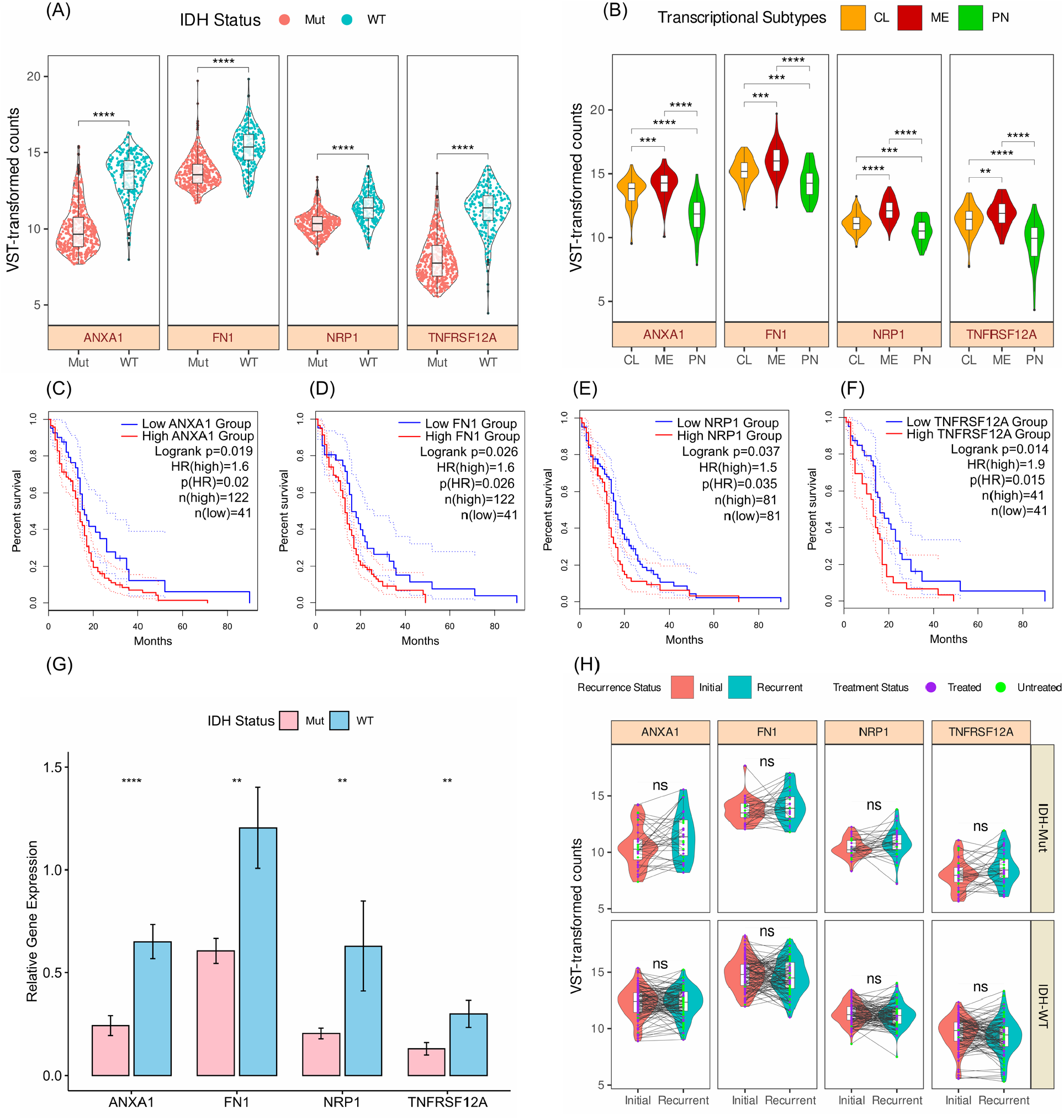
Expression analysis of *ANXA1, FN1, NRP1* and *TNFRSF12A* genes in bulk IDH1-stratified samples. (A) Boxplots with violins depict the expression levels of *ANXA1, FN1, NRP1*, and *TNFRSF12A* genes in TCGA-GBM and TCGA-LGG cohorts, stratified by *IDH1* mutation status. Boxes represent the interquartile range (IQR), with the median indicated by the center line. Whiskers extend to the minimum and maximum values within 1.5 times the IQR. Violins depict the distribution of expression levels for each group; (B) Expression levels of candidate genes in CL (n = 87), ME (n = 77), PN (n = 42) transcriptomic subtypes of TCGA-GBM; (C-F) Kaplan-Meier survival curves illustrating the prognostic significance of *ANXA1, FN1, NRP1*, and *TNFRSF12A* genes in overall survival among TCGA-GBM cohort; (G) Bar plots depict the relative gene expression levels of represented candidate genes in IDH-WT (n = 39) and IDH-Mut (n = 41) samples as determined by RT-qPCR; (H) longitudinal expression analysis of candidate genes in paired primary and recurrent samples using GLASS bulk RNA-seq. P-values were determined using two-sided Wilcoxon rank-sum (A, B, G) and signed-rank test (H). Results were considered statistically significant when *p < 0.05, **p < 0.001, ***p < 0.0001, ****p < 0.00001. ns indicates no significant difference.

In follow-up, we used the Glioma Longitudinal AnalySiS (GLASS) bulk RNA-seq dataset to evaluate expression of *ANXA1, FN1, NRP1* and *TNFRSF12A* genes between paired initial and recurrent samples in IDH-WT (n = 124) and IDH-Mut (n = 43) patients. The almost invariable recurrence of GBM despite initial treatments with surgery and chemoradiation, poses a greater challenge as there is no standard of care at disease progression. Recent large scale longitudinal efforts for uncovering unique evolutionary properties that primary tumors acquire after recurrence have yet concluded with a lack of selective pressure against specific genetic alterations [74]. New therapeutic insights could emerge from studying the interactive landscape of glioma TME over the course of disease progression. We observed no significant difference in expression of the candidate genes between initial and recurrent cases of either IDH-WT or IDH-mutant gliomas (Fig. 6H; Table S10, S11).

## DISCUSSION

A growing body of evidence underscores pivotal roles of TAMs in various aspects of glioma biology. In order to investigate whether adverse prognosis of IDH-WT gliomas compared to IDH-Mut counterparts can be attributed, at least in part, to their differences in TME architecture and communications, we conducted a comprehensive analysis. First, we provided a high-resolution single-cell view of the myeloid compartment within the glioma TME, delineating BMD, MG-TAMs into seven functional transcriptomic clusters. Furthermore, we compared different cell type composition patterns in IDH-WT and IDH-Mut. Of note, two clusters with mesenchymal phenotype were reminiscent of macrophages that mimic phenotypes of tumor cells and were differentially present in the IDH-WT group.

Our study revealed specific myeloid subsets highly associated with alterations in signature genes commonly mutated in glioma, achieved by integrating bulk expression profiles and phenotypic data from TCGA-GBM and TCGA-LGG cohorts. Our analysis underscored the significant dependence of the glioma immunophenotype on tumor genotype, providing further evidence that genetic alterations in cancer cells shape the immune landscape of tumors [75]. These results offer insights to tailor personalized combination therapies that target both myeloid and cancer cells, leveraging the underlying genetic profile. Utilizing survival metadata available through the GDC portal, we also investigated the association between myeloid clusters and overall survival of GBM patients. Two BMD-TAM clusters with mesenchymal phenotype were highly linked to worse overall survival. By analyzing transcriptomic data from a bulk RNA-seq dataset of GBM patients treated with anti-PD-1 antibodies, we also noted enrichment of the mesenchymal BMD-TAM signature in the non-responder group, indicating association of these macrophages with efficacy of ICI therapy. We also observed that mesenchymal BMD-TAMs exhibit lower expression levels of the *CSF1R* gene (Fig. S2A), which could potentially diminish efficacy of CSFR inhibitors in targeting these cells. This observation may explain the failure of CSF1R inhibitors in clinical trials on recurrent GBM [76]. Our findings suggest that SLAMF9 may be a promising target for mesenchymal BMD-TAMs in GBM. Future functional studies are warranted to validate this hypothesis and elucidate the underlying mechanisms.

This work also provided mechanistic insights into the regulatory landscape of glioma TAMs accompanied by their trajectory paths and gene expression dynamics. Our analysis, which particularly investigated two mesenchymal clusters within BMD-TAMs, identified PPARG and ELK1 in BMD-TAM.Mes and MXI1 in BMD-TAM.Mes/Hypo as potential master TFs. PPARγ plays a crucial role in terminal differentiation of MMP9^+^ TAMs In hepatocellular carcinoma (HCC), with knockdown of corresponding gene or treatment with PPARγ inhibitors resulting in a significant decrease in MMP+ macrophages [77]. Series of *in vivo* and *in vitro* experiments are required to fully study the identified TFs in mesenchymal BMD-TAMs of GBM.

Using scRNA-seq, we inferred a cell-cell interaction network that revealed both the established and the less characterized tumor-TAM crosstalk. By comparing the inferred interactome between IDH-WT and IDH-Mut tumors, we learned that unique communication of GBM with TAMs primarily stems from its MES-like state, which is absent in IDH-Mut tumors. This observation implies a therapeutic strategy: shifting malignant states towards MES-like states, followed by targeted elimination, could potentially hamper tumor-supportive networks. By integrating scRNA-seq with ST, we demonstrated the spatial significance of four L-R interactions linked to the distinguishing phenotypes of GBM tumors. According to our findings, ANXA1 ligand, derived from MES-like malignant states of IDH-WT GBM, interacts with FPR1/3 receptors on BMD-TAMs and MG-TAMs. In a study by Wu and colleagues [55], *ANXA1* knockdown was found to significantly reduce macrophage-mediated suppression of CD8+ T cells and tumor progression in GBM mouse models. Additionally, in a recent study, Zheng *et al*. [64] demonstrated that decreasing ANXA1 expression improves response to Toll-like receptor 3 (TLR3) ligands and that ANXA1 has the potential to serve as a reliable predictive marker of patient response to TLR agonists. Taken together, ANXA1 poses an obstacle in eliciting an effective immune response in GBM as MES-like cells instruct an immunosuppressive TME through the ANXA1-FPR1 axis. We also demonstrated significant FN1-integrin interactions between FN1^high^ BMD-TAM.Mes and tumor cells in hypoxic areas. Hypoxia can trigger the regulatory signaling pathways of EMT [78]. In addition to guiding the necessary orientation for invasion, FN1-induced signaling seems to prime cells for extensive spreading [79]. In a recent study on gastric cancer, researchers demonstrated that blocking the integrin complex could prevent metastasis promoted by FN1^high^ TAMs via FN1-integrin interactions [78]. In another study on HCC, FN1 derived from TAMs and fibroblasts facilitated metastasis and modulated transcriptomic subtypes through the JUN-Hippo signaling pathway [80]. Collectively, our analysis indicates that FN1^high^ BMD-TAM.Mes cells enhance tumor cell migration and invasion by providing fibronectins as adhesive substrates and signaling cues. Additionally, we reported the VEGFA-NRP1 signaling axis between MES-like cells and BMD-TAMs, particularly enriched in hypoxic regions of the TME. Disruption of NRP1 signaling in macrophages, as demonstrated by Casazza *et al*. [67], either through depletion or targeted blockade, significantly impedes tumor progression. This effect is amplified by preventing TAM infiltration into hypoxic niches. Preclinical data highlights NRP1 antagonism as a promising anti-tumor strategy, acting through a dual mechanism: inhibiting tumor angiogenesis and directly suppressing cancer cell survival or proliferation, potentially effective across diverse malignancies [67, 81-83]. In this context, NRP1 on the surface of TAMs likely promotes their migration towards angiogenic areas, thereby facilitating their pro-tumoral functions. We also demonstrated TWEAK ligands secreted by MG-TAMs, combine with their Fn14 receptors largely on MES-like cells, highlighting the importance of MG-TAMs in GBM pathogenesis. Targeted therapy against TWEAK/Fn14 or disruption of downstream noncanonical NF-κB signaling via inhibition of NF-κB-inducing kinase (NIK) has been suggested as therapeutic strategies against GBM [72, 84].

We finally, demonstrated elevated mRNA levels of *ANXA1, FN1, NRP1*, and *TNFRSF12A* genes in IDH-WT compared to IDH-Mut gliomas using human tissue samples. Bulk RNA-seq analysis further revealed that the GBM-ME subtype exhibits the highest expression levels for the candidate genes. These observations align with the established dominance of the MES-like state in IDH-WT gliomas, which is characterized by its reliance on the TME [16]. The longitudinal expression follow-up, on the other hand, demonstrated no significant difference between paired primary and recurrent cases in either IDH-WT or IDH-Mut categories, indicating that the interactive network of malignant states in glioma with their associated TAMs may not undergo drastic changes upon recurrence. A recent study by Hoogstrate *et al*. [85] showed that longitudinal evolution of IDH-WT GBM is shaped predominantly by re-organization of the TME and in corroboration of previous findings about transcriptional shifts towards MES-like phenotype adds that this preferred evolutionary path may be steered by the TME and is often observed upon recurrence. In essence, these findings suggest that CL subtypes remain more stable than both ME and PN subtypes and constitute a larger proportion of recurrent tumors [11, 85]. Furthermore, all three phenotypes associated with recurrence – neuronal, mesenchymal and proliferative – have been observed in both IDH-defined categories of glioma, albeit with varying frequencies. IDH-WT tumors tend to exhibit all three phenotypes upon recurrence, whereas only a subset of IDH-Mut tumors display the proliferative phenotype [86]. Moreover, a recent longitudinal investigation reports that most IDH-WT and IDH-Mut tumors retain their original subtype upon recurrence, with only a small fraction switching subtypes. The study also suggests that standard therapies may contribute to loss of DNA methylation in IDH-Mut gliomas. This demethylation event could lead to the epigenetic activation of genes associated with tumor progression, creating a TME resembling treatment-naive IDH-WT gliomas [87]. Taken together, these findings might provide an explanation for our observations.

## CONCLUSIONS

Overall, focusing on the myeloid compartment in IDH-stratified diffuse gliomas, we demonstrate that the GBM-trained BMD-TAMs with mesenchymal phenotype can be harnessed for novel immunotherapeutic strategies. Our study on characterization of the bidirectional interactome of tumor cells and TAMs, also provides important insights into the GBM-unique sources for immunosuppression, proliferation and metastasis that can be tailored for future investigations.

## Acknowledgments

We thank the patients and their families for their generous contributions to this research and the research community. We also thank the staff at the neurosurgery and pathology departments of Shariati hospital for their support during sample collection.

## Author contributions

MM, MD and MT conceptualized and designed the study. MA, SB, AK and MZ provided the glioma tissue samples. HS classified the samples according to histopathological findings. MM and MD did the experiments, analyzed the data and prepared the figures. AN, SB, AM assisted with the experiments. MM and MD wrote the first draft of the manuscript. MT supervised and reviewed the work.

## Data accessibility

The scRNA-seq dataset generated by [88] study was obtained from the Synapse portal (https://synapse.org/singlecellglioma). Spatial transcriptomic datasets from [89] study was downloaded from Datadryad (https://doi.org/10.5061/dryad.h70rxwdmj). The bulk RNA-seq datasets supporting the findings in the current paper are publicly available from the TCGA database (https://portal.gdc.cancer.gov/, via TCGAbiolinks v. 3.19 [90]), Synapse portal (https://www.synapse.org/#!Synapse:syn17038081/wiki/585622) for GLASS bulk RNA-seq and PRJNA482620 (https://www.ncbi.nlm.nih.gov/bioproject/PRJNA482620/) for Zhao et al. The IVY-GAP data including the bulk RNA-seq and RNA-ISH are publicly available at https://glioblastoma.alleninstitute.org/.

## Supplementary data

**Fig S1**. Characteristics and annotation of glioma scRNA-seq dataset.

**Fig S2**. Expression patterns of *CSF1R* and *SLAMF9* genes in glioma myeloid cells and in normal human brain.

**Fig S3**. Estimation of cellular fate across BMD-TAMs via CellRank.

**Fig S4**. Expression analysis of *FPR1, FPR3, NRP1, NRP2, FN1* and *TNFSF12* genes across myeloid compartment.

**Table S1**. List of gene signatures used for scRNA-seq analysis.

**Table S2**. Sample information.

**Table S3**. List of primers used in the study.

**Table S4**. Scaled AUC scores of gene signatures in glioma myeloid cells.

**Table S5**. List of differentially expressed genes in glioma myeloid clusters.

**Table S6**. List of differentially expressed genes between IDH-WT and IDH-Mut tumors per BMD-TAM.

**Table S7**. List of differentially expressed genes between IDH-WT and IDH-Mut tumors per MG-TAM.

**Table S8**. mesenchymal BMD-TAM gene signature.

**Table S9**. List of differentially expressed genes between IDH-WT and IDH-Mut bulk RNA-seq cases.

**Table S10**. List of differentially expressed genes between IDH-WT paired initial and recurrent GLASS bulk RNA-seq.

**Table S11**. List of differentially expressed genes between IDH-Mut paired initial and recurrent GLASS bulk RNA-seq.

