## Supplementary Figures for "Distinct Tumor-TAM Interactions in IDH-Stratified Glioma Microenvironments unveiled by Single-Cell and Spatial Transcriptomics"

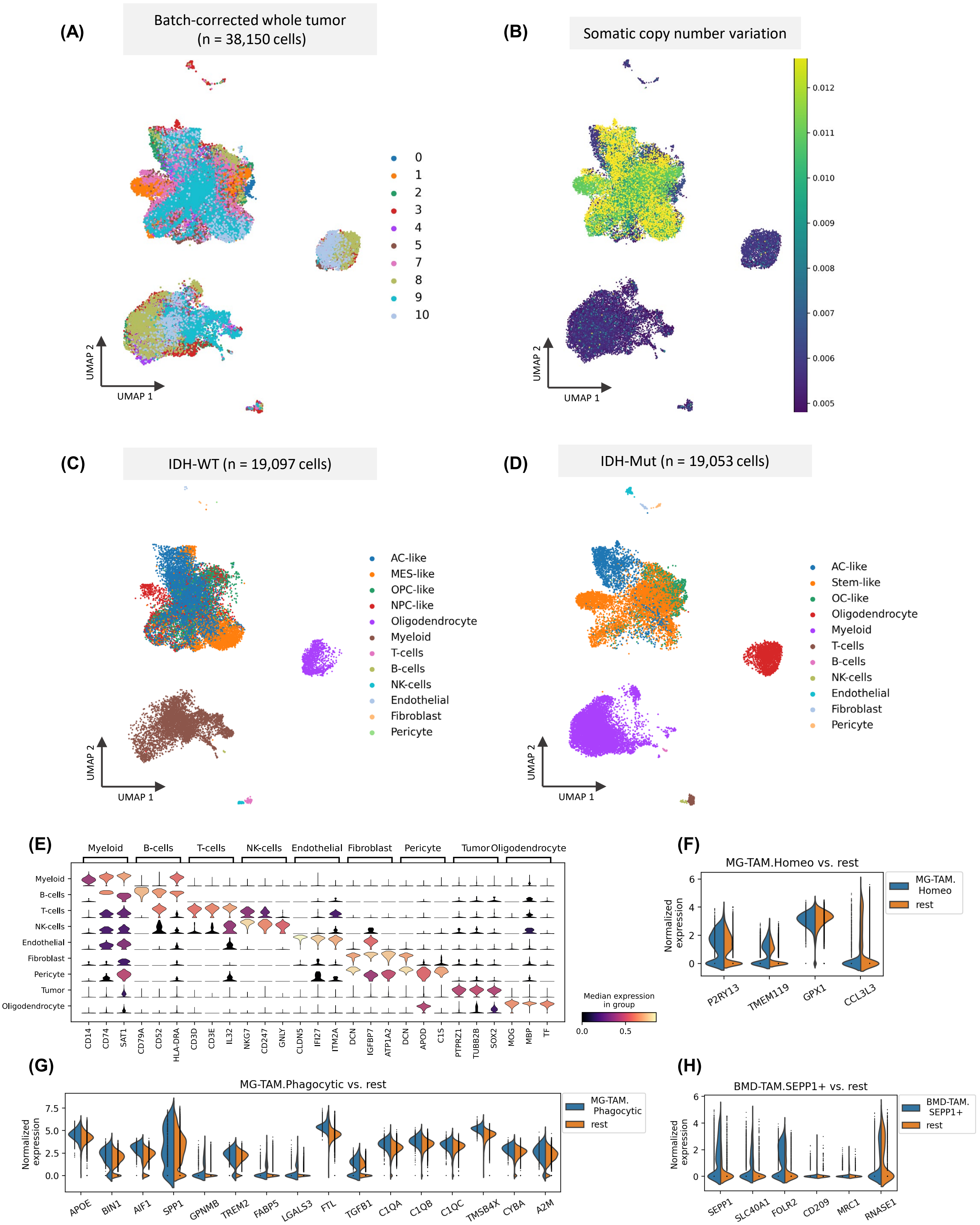


Fig. S1. Preprocessing and annotation of glioma scRNA-seq dataset. (A) UMAP plot showing the batch corrected scRNA-seq dataset. Each dot depicts a single cell and colors refer to the sample of origin; (B) UMAP plot visualizing the inferred score of SCNA across scRNA-seq dataset; (C) UMAP visualization of scRNA-seq from IDH-WT patients. Colors refer to the annotated cell types; (D) UMAP visualization of scRNA-seq from IDH-Mut patients. Colors refer to the annotated cell types; (E) Stacked violin plots showing the average gene expression levels of identified biomarkers for major clusters presented in Fig. 1A (left); (F-H) Split violin plots exhibiting the expression intensity distribution of represented marker genes in (F) MG-TAM.Homeo, (G) MG-TAM.Phagocytic cells and (H) BMD-TAM.SEPP1+ vs. the rest of myeloid compartment.

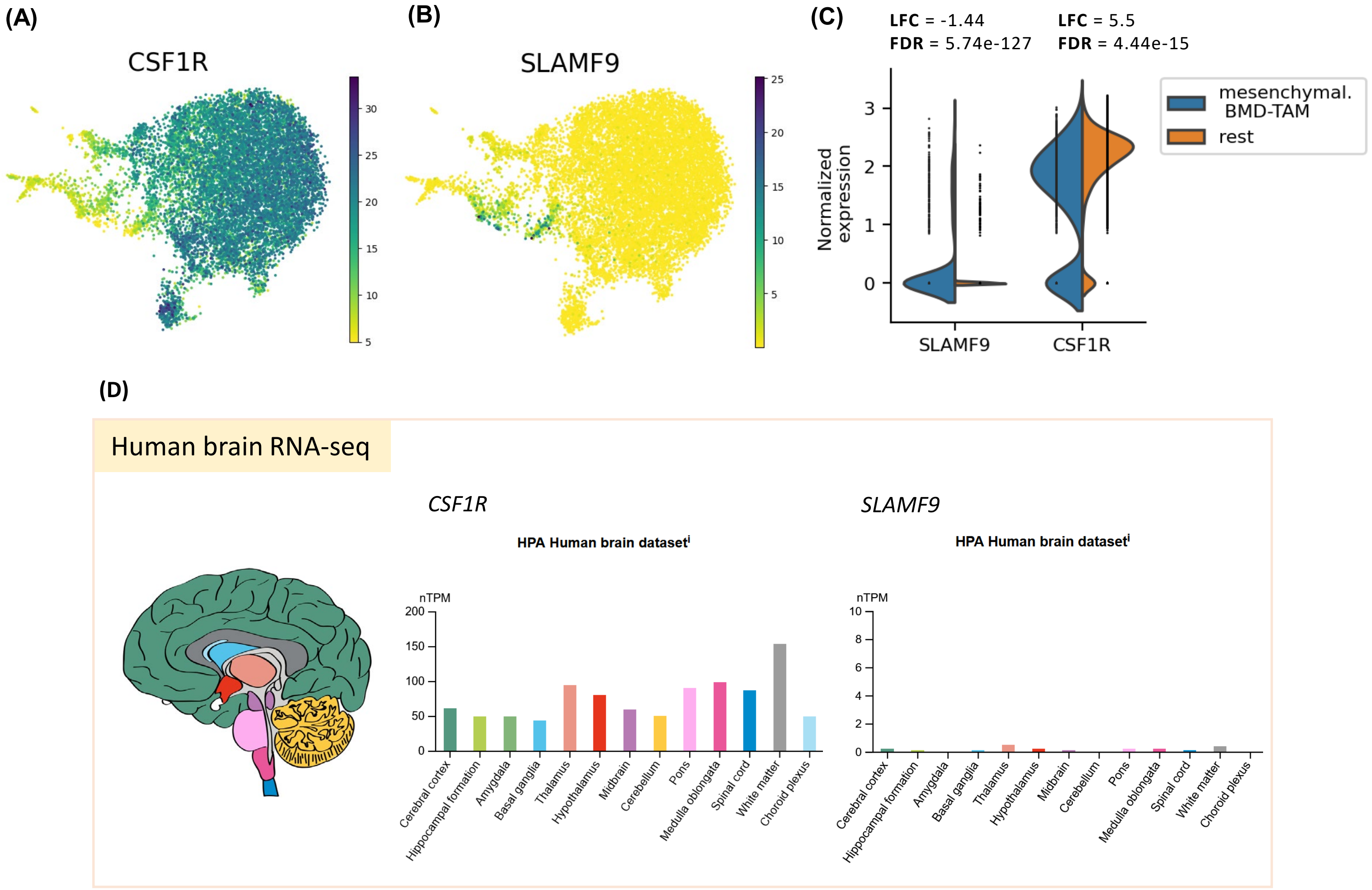


Fig. S2. Expression patterns of *CSF1R* and *SLAMF9* genes in glioma myeloid cells and in normal human brain. (A) UMAP plot showing normalized expression levels of *CSF1R* gene in glioma myeloid compartment; (B) UMAP plot showing normalized expression levels of *SLAMF9* gene in glioma myeloid compartment; (C) Split violin plot comparing the expression of *CSF1R* and *SLAMF9* genes between mesenchymal BMD-TAMs and rest of the myeloid compartment; (D) Bar plots showing expression levels of *CSF1R* and *SLAMF9* genes in bulk RNA-seq datasets from different anatomical regions of normal human brain (www.proteinatlas.org). LFC, Log Fold Change; FDR, False Discovery Rate.

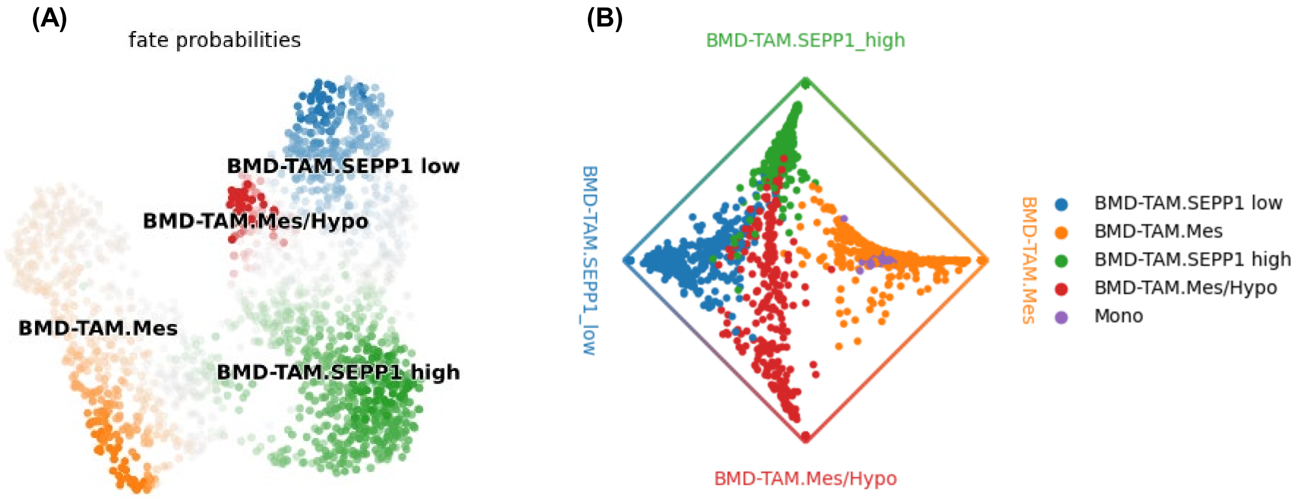


Fig. S3. CellRank analysis estimates cellular fate bias in BMD-TAM states. (A) UMAP embedding visualization of CellRank's predicted fate probabilities towards identified cellular macrostates; Cells are colored according to their most likely terminal cell state, with color intensity reflecting the strength of the lineage bias; (B) Circular projection of cells colored by cluster annotations, reflecting fate probabilities towards the identified macrostates.

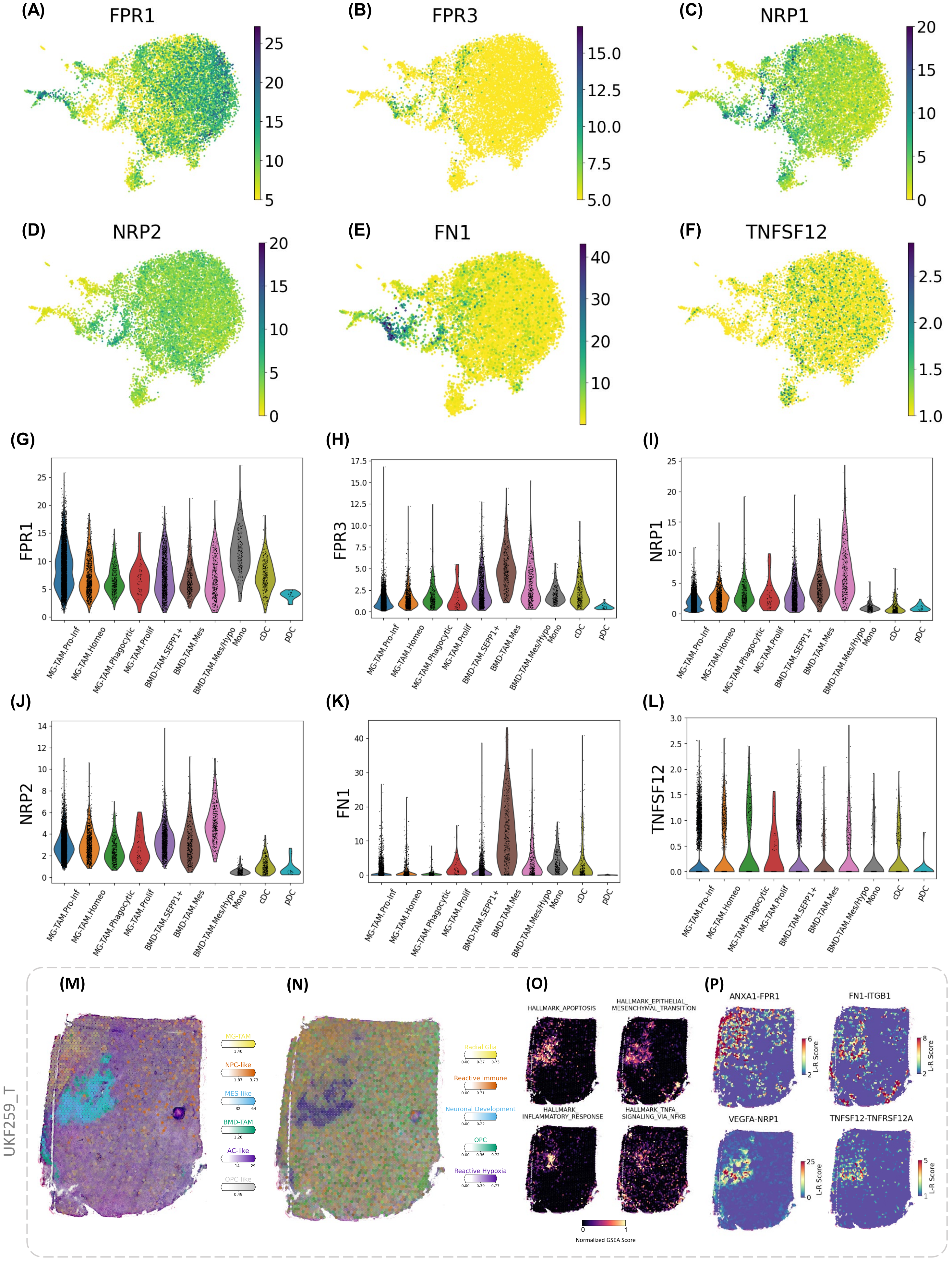


Fig. S4. Expression analysis of highlighted genes across myeloid compartment and complementary analysis of ST. (A-F) UMAP plots demonstrating the expression of *FPR1, FPR3, NRP1, NRP2, FN1* and *TNFSF12* genes in glioma myeloid compartment; (G-L) violin plots demonstrating the expression of *FPR1, FPR3, NRP1, NRP2, FN1* and *TNFSF12* genes in myeloid clusters; (M) Surface plot showing the spatial locations of cell types in Visium ST datasets from UKF259_T donor; (N) Surface plot indicating spatial enrichment scores for five transcriptional programs; (O) Surface plots showing spatial GSEA scores for MSigDb hallmark gene sets; (P) Spatial cell-cell interaction of representative L-R pairs.
